# Integrated single-cell potency and expression landscape in mammary epithelium reveals novel bipotent-like cells associated with breast cancer risk

**DOI:** 10.1101/496471

**Authors:** Andrew E. Teschendorff, Samuel J Morabito, Kai Kessenbrock, Kerstin Meyer

## Abstract

The identification of progenitor and stem like cells in epithelial tissues, as well as those that may serve as the cell of origin for epithelial cancers, is an outstanding challenge. Here we present a novel algorithm, called LandSCENT, which constructs a 3-dimensional integrated landscape of cell-states, encompassing cell-potency and expression subtypes, to facilitate the identification of progenitor and stem-like cells. Application to thousands of single-cell RNA-Seq profiles from the normal mammary epithelium reveals a rare 5% subpopulation of highly potent single-cells. The integrated landscape naturally predicts that these cells define a bi-potent-like state, a result not obtainable via standard methods or without invoking prior assumptions. The bi-potent-like cells are overrepresented within the basal compartment but also overlap with an immature luminal phenotype. We characterize the transcriptome of these cells and show that is enriched for a mammary stem-cell module. We further identify *YBX1*, a regulator of breast cancer risk identified from GWAS, as the key transcription factor defining this candidate bi-potent cellular phenotype. We validate the putative bi-potency of *YBX1*-marked cells using independent FACS-sorted bulk expression data. In addition, *YBX1* is overexpressed in basal breast cancer and correlates with clinical outcome. In summary, we here provide a novel computational framework which may serve to identify and prioritize candidate normal or cancer progenitor/stem-like single-cell phenotypes, for subsequent functional studies.

## Introduction

Single-cell RNA-sequencing (scRNA-Seq) studies are revolutionizing our understanding of cellular development, helping us to elucidate the hierarchical organization of cell-types within complex tissues (Patel et al. 2014; Trapnell et al. 2014; Treutlein et al. 2014; Scialdone et al. 2016; Tirosh et al. 2016a; Tirosh et al. 2016b; Treutlein et al. 2016; Haber et al. 2017; Regev et al. 2017; Rozenblatt-Rosen et al. 2017; Hon et al. 2018; Laurenti and Gottgens 2018; Shepherd et al. 2018). In these studies, a common computational task is the clustering of single cells, which may reveal novel cell-types within these tissues (Haber et al. 2017; Han et al. 2018). Relations between known and novel cell-types can be subsequently derived using lineage-trajectory inference type algorithms (Trapnell et al. 2014) (Chen et al. 2016) (Marco et al. 2014) (Haghverdi et al. 2016) (Grun et al. 2016). However, assignment of single cells to cell-types often requires prior knowledge of specific markers, which may inevitably introduce bias (Stingl et al. 2006; Trapnell 2015; Yuan et al. 2017). In certain circumstances this bias can be substantial, specially if knowledge of suitable markers is not available or at best controversial (Costa et al. 2018; Grun 2018). Moreover, lineage-trajectory inference algorithms, including recent state-of-the-art ones such as Monocle-2 (Qiu et al. 2017), often require specification of a “root” cell or node, in order to give the trajectories a “temporal” direction. In the absence of temporal data, the specification of this root node may rely on existing biological knowledge and therefore equally subject to bias. Another related and key problem is that cell-types are typically inferred as clusters of relatively high cell-density in a two dimensional reduced space, a procedure which does not necessarily allow for the identification of cellular states. For instance, how to identify novel progenitor or stem-like states within a cell-type may not be possible using two-dimensional clustering alone since potency/stemness may be defined by additional latent dimensions.

To address these outstanding challenges, we here present LandSCENT, a novel computational framework that avoids the aforementioned biases, assigning each cell, not only to a specific cell-type, but also to a specific potency state (Teschendorff and Enver 2017). LandSCENT achieves this without the need for prior knowledge or assumptions. LandSCENT integrates the inferred cell-types and potency states into a multi-layered single-cell landscape, where cell-states are defined by clusters of single-cells within a potency state. This novel approach allows cells to be placed into specific cellular-states, thus allowing novel cellular phenotypes to be identified, for instance novel progenitor or stem-like states within complex epithelial tissues.

We illustrate this strategy in the context of the breast epithelium, a tissue for which scRNA-Seq encompassing over 25,000 single epithelial cells from 4 women, has recently been generated using the 10X Genomics Chromium assay (Nguyen et al. 2018). We apply LandSCENT to this data to construct an integrated potency and cell-type landscape at the single-cell level. This landscape reveals a novel putative bi-potent progenitor like cell-state, characterized by overexpression of *YBX1*, a recently discovered regulator of breast cancer risk (Castro et al. 2016), a result which we would not have found had we used standard state-of-the-art clustering methods. We further validate the bi-potent/stem-like nature of the identified single-cells using orthogonal bulk expression data of mammosphere-derived mammary stem cells. Our data support the view that the identified bi-potent cells expressing *YBX1* may give rise to both basal and luminal progenitors, potentially marking the cell-of-origin for basal breast cancer.

## Results

### Constructing an integrative landscape of cell-states in breast epithelium

We posited that improved clustering of single-cells so as to reveal novel biology, would be possible by integrating cellular state information with the cells’ expression profiles when performing the clustering itself. One important feature of a cell that informs on cell-state is its differentiation potency and in previous studies we proposed and validated an *in-silico* measure of single-cell potency, based on the concept of single cell signaling entropy (SCENT) (Banerji et al. 2013; Teschendorff and Enver 2017), which we have further shown is more robust than other proposed single-cell potency models (Grun et al. 2016; Guo et al. 2016; Shi et al. 2018a). We stress that SCENT represents a marker-free systems-biology approach to the quantification of a cell’s potency, which has been demonstrated to be very robust, and which is applicable also to bulk samples (Banerji et al. 2013; Teschendorff and Enver 2017; Shi et al. 2018b). This is important because the alternative approach, i.e. to use expression of surface markers, is unlikely to capture the full biological complexity underlying cellular potency, while also introducing potential bias. Thus, here we present LandSCENT, a novel extension of SCENT that combines inference of cell potency with single-cell clustering to construct a landscape of single-cell states: these single-cell states integrate the single-cell potency estimates with the inferred cell-type clusters, providing a 3-dimensional landscape representation (**Fig.1**, **Methods**). Here we applied LandSCENT to a 10X Genomics Chromium assay profiling thousands of single-cells in the breast epithelium (Nguyen et al. 2018), in order to define the landscape of cellular states in this tissue (**Fig.1**).

**Figure-1:**
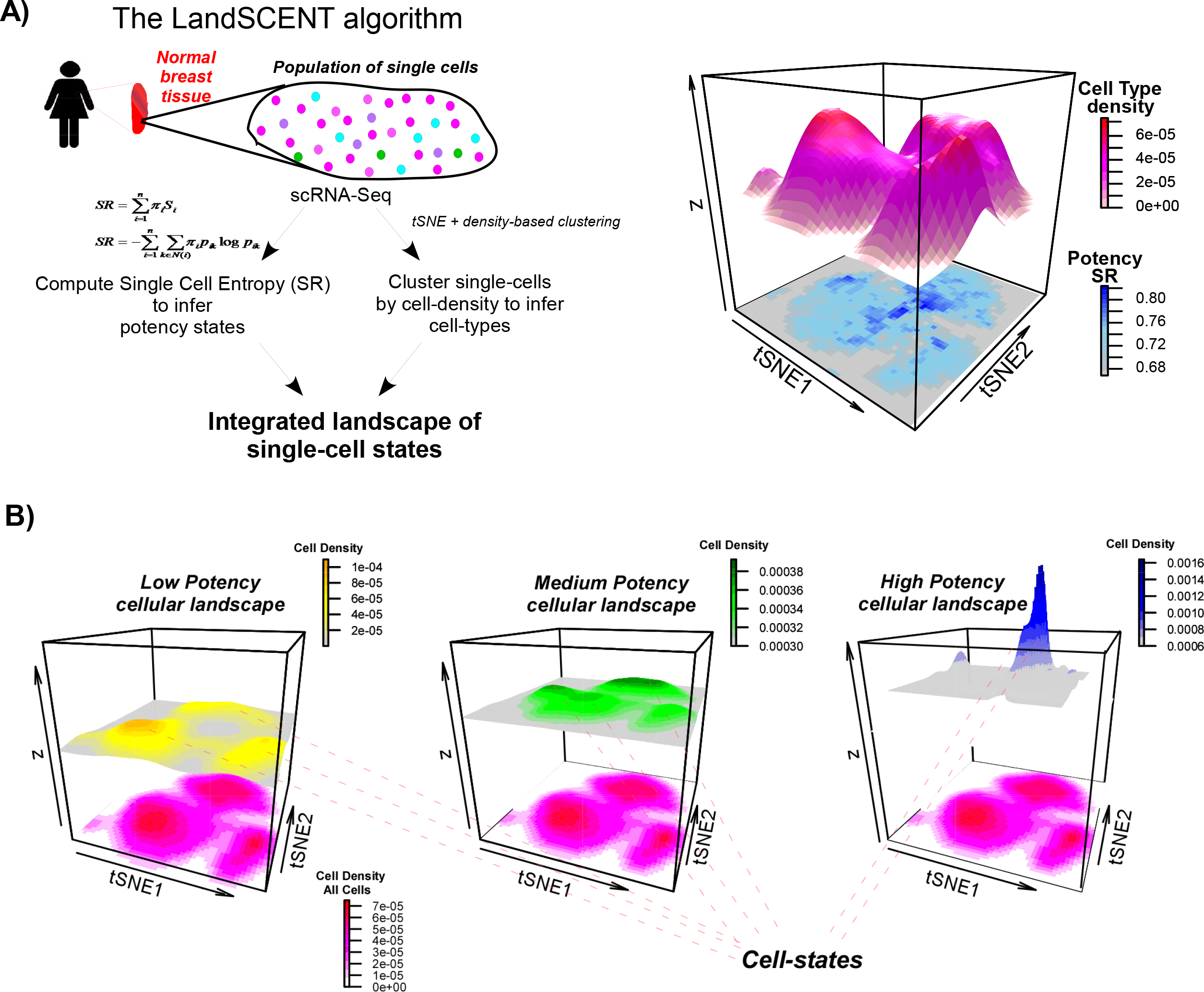
Flowchart of the LandSCENT algorithm to construct an integrative landscape of cell-states from scRNA-Seq data. **A)** Left: Signaling entropy (SR) is applied to the scRNA-Seq profile of each individual cell to estimate its differentiation potency and to infer potency states. Clustering of single-cells is performed with t-SNE followed by density based spatial clustering to identify clusters of high cell-density, which we call cell-types. Right: Surface cell-density map representation in t-SNE space for all single cells, showing the main cell-types, with the smoothed SR (potency) values projected at the bottom. **B)**An example of an integrated layered landscape of cellular states, where surface cell-density maps are shown for cells in each inferred potency state (low, medium and high potency), defining cell-states within or between major cell-types. The integrated landscape can reveal cell-states not discernable via standard two dimensional clustering (shown at the bottom of each landscape).

First, we phenotypically characterized the single cells, by performing t-SNE (van der Maaten 2008) followed by density-based spatial clustering (Ester et al. 1996) on 3473 single epithelial cells (after QC) from one individual and using a reduced subset of 4261 genes that exhibited a significant average and variance in expression across all cells (**Methods**). This revealed three main single-cell clusters (**Fig.2A**), in line with previous observations (Nguyen et al. 2018). One of these clusters expressed high levels of *KRT14*, a well-known basal marker, which was not expressed in the other two main clusters (**Fig.2B**). Instead, the other two clusters expressed *KRT18*, a well-known luminal marker. Consistent with the report of Nguyen et al (Nguyen et al. 2018), the two luminal clusters were distinguished by expression of lactotransferin (*LTF*) and luminal differentiation markers (*GATA3/FOXA1*), as well as hormone receptors (*ESR1/PGR*) (**Fig.2B**), suggesting that the higher *LTF*-expressing cluster represents a more immature (alveolar) luminal phenotype.

Next, we applied our Signaling Entropy Rate (SR) measure from SCENT to estimate the differentiation potency of each single cell. To broadly categorize different levels of inferred potency, we applied a Gaussian mixture model to the logit-transformed potency estimates of the 3473 single cells, revealing the existence of three main potency states (**Fig.2C-D**, **Methods**). We observed that the highest potency state represented a minority population, with approximately only 5% of single-cells falling into this putative progenitor or stem-like state (**Fig.2D**).

**Figure-2:**
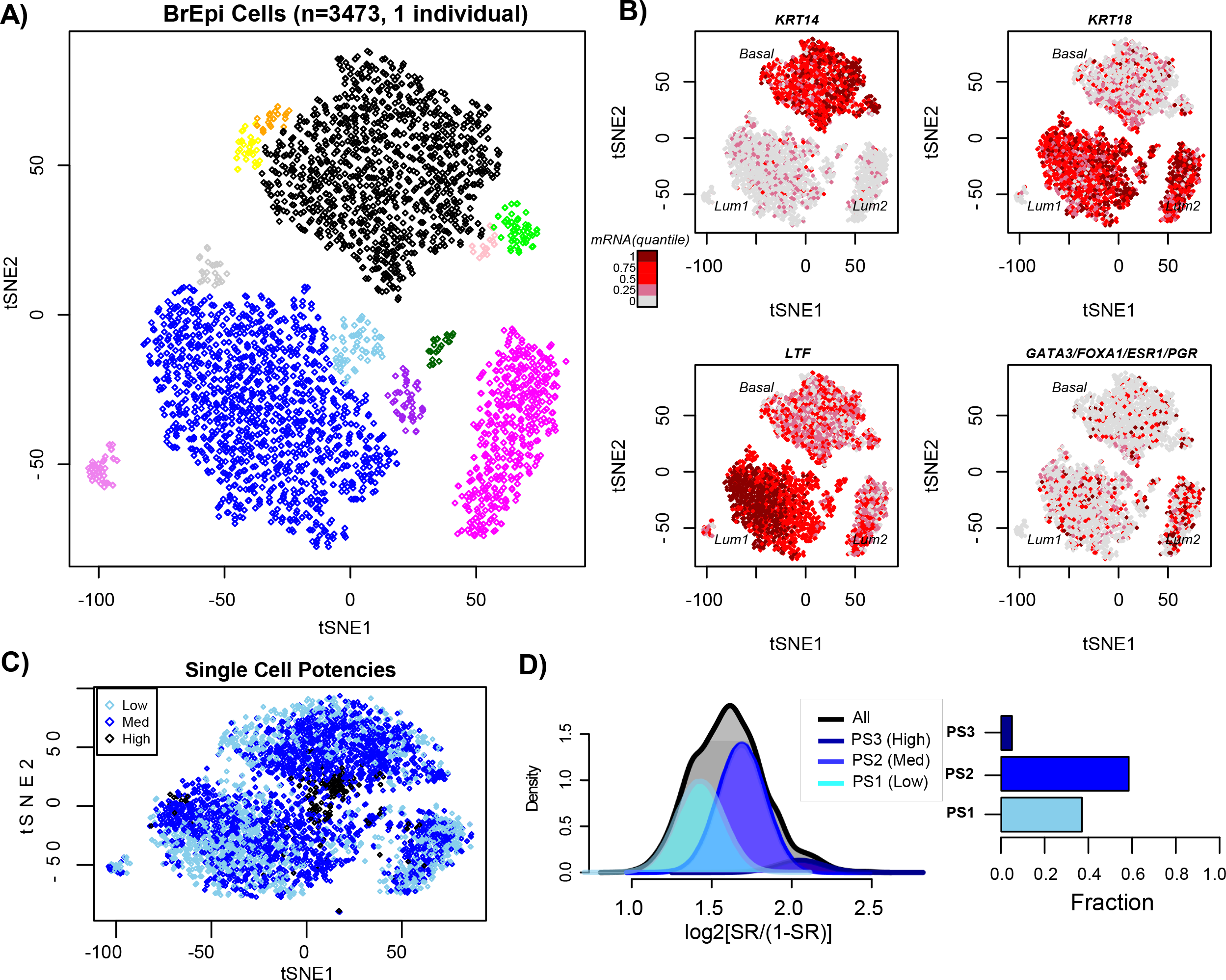
Inferring cell-types and potency states in breast epithelium. **A)** t-SNE clustering diagram for single-cells derived from one individual (Ind-4). Single-clusters were inferred with dbscan and are labeled with different colors. Of note, single-cells that mapped to the periphery of clusters and therefore were not assigned to any cluster have been suppressed. **B)** As A), but now with the single cells labeled by expression levels of *KRT14* (a basal marker), *KRT18* (a luminal marker), *LTF* (lactotransferin) and mean expression of *GATA3, FOXA1, ESR1* and *PGR*, as indicated. Different quantiles of expression levels of each marker are indicated by color with brown indicating high expression and grey low expression. **C)** As A), but now displaying all single cells (i.e. including those mapping to the periphery of clusters) and with single-cells labeled by the inferred potency state (see D)). **D) Left panel:**Gaussian mixture model fit to the logit transformed SR values (x-axis) from 3473 single cells infers 3 potency states. The density distributions for all cells (black line) and those for the inferred mixture components (different shades of blue) are shown. The Bayesian Information Criterion (BIC) was used to select the optimal number of potency states, which in this case was found to be 3 (PS1, PS2, PS3). **Right panel:** Percentage barplot indicating the fraction of single-cells assigned to each of the three potency states.

### Validation of potency assignments

Although signaling entropy has been extensively validated as a cell-potency measure (Teschendorff and Enver 2017; Shi et al. 2018a), we sought additional validation of the specific potency assignments in the current dataset. It is well known that *GATA3, FOXA1* and *ESR1* are associated with a more differentiated luminal phenotype and therefore the expectation would be that their expression levels should be higher in the luminal cells of lowest potency. We were able to confirm this with high statistical significance (**Fig.3A**). We also validated the potency assignments within the basal compartment. For instance, we observed that expression of *KRT5* and *EGFR*, two well-known basal differentiation markers, decreased in the basal cells of higher potency (**Fig.3B**). We note that all these negative correlations were apparent only when we restricted to cells where the genes were expressed. If all cells were included, including technical and biological dropouts, we did not observe these genes to exhibit the expected negative correlation: in fact, they showed an opposite trend due to a larger number of dropouts among low potency cells (**SI fig.S1**). To investigate this further and to validate our method to call differential expression (DE), we used bulk mRNA expression data from FACS sorted differentiated luminal and basal cells (Shehata et al. 2012) to define a gold-standard list of 5,773 differentially expressed genes between basal and luminal cells. For each of these gold-standard genes, and using only cells expressing the corresponding gene, we derived a t-statistic of differential expression between the single cell basal and luminal clusters, which revealed that for the great majority of gold-standard genes with sufficient single-cell data, these exhibited the expected pattern of differential expression (OR=8.31, Fisher=test P=2e-26, **SI fig.S2**). Based on this, we conclude that performing DE using only cells expressing the gene is a valid procedure, thus also validating our potency assignments.

**Figure-3:**
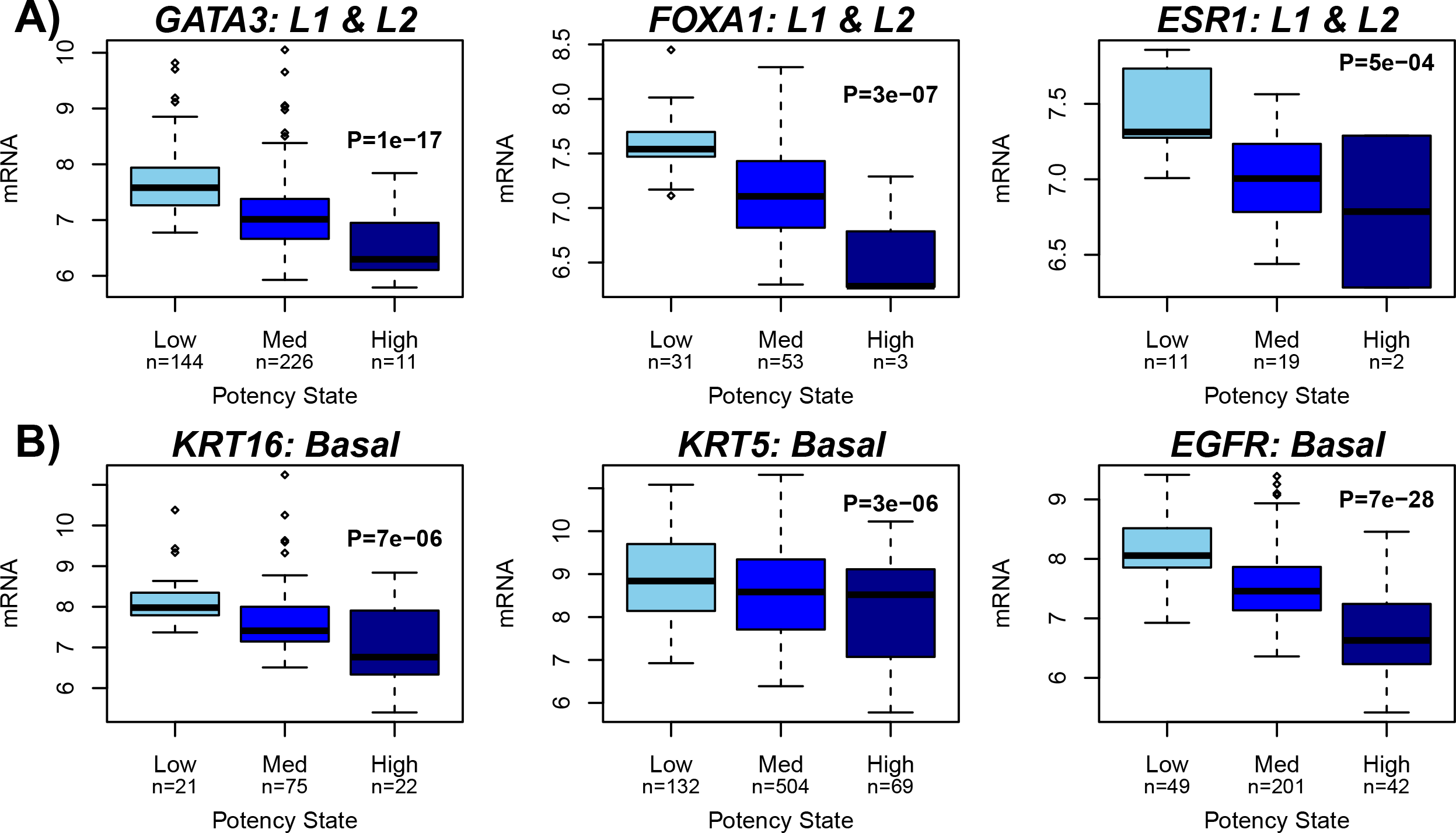
Validation of potency assignments. **A)** Boxplots of normalized log-expression (y-axis) for known markers of luminal differentiated cells (*GATA3, FOXA1*) and hormone receptor (*ESR1*) against inferred potency state (x-axis) for all single cells assigned to the two main luminal clusters (L1 & L2) and further restricting to cells where these genes are expressed. Numbers of single-cells assigned to each potency state is given. P-value is from a (two-tailed) linear regression. **B)** As A), but for known basal differentiation markers (*KRT16, KRT5, EGFR*) and restricting to cells that were assigned to the basal cluster.

### Integrative landscape reveals a putative bi-potent cell state

Having identified and validated the main single-cell clusters and potency states, we next considered the distribution of potency states across these 3 clusters, as well as those cells not assigned to any cluster (“peripheral cells”). Interestingly, cells in the high potency state were found primarily within the basal compartment, but also mapped preferentially to the common peripheral area of the three main clusters, and were therefore also relatively over represented among peripheral cells (**Fig.2C**, **Fig.4A**). To assess this in more detail, we used LandSCENT to create cell-density elevation maps of all cells, and separately also for all highly potent cells, within the two-dimensional t-SNE landscape, which confirmed that the maximum density of the highly potent cells defined a peak within the basal cluster, but with a ridge connecting it to another peak within the immature luminal (L1) cluster (**Fig.4B**), suggestive of a bi-potent cell population. In line with this, we observed that among all cells categorized into the high potency (PS3) state, those falling within this peak also exhibited the highest levels of signaling entropy (**SI fig.S3**). To exclude the possibility that these higher or bi-potent cells may be doublets, we estimated doublet scores for all cells using a novel simulation approach (Dahlin et al. 2018). In line with the expected doublet rate for 10X technology, this analysis revealed that approximately 2% of the assayed cells are potential doublets (**SI fig.S4A**). As expected, most of these mapped to the peripheral area between the major luminal and basal clusters, yet they clearly also did not overlap with the most highly potent cells within the basal and luminal clusters, confirming that our candidate bi-potent cells are generally not doublets (**SI fig.4B**). Supporting this, we observed that the relation between signaling entropy and doublet scores is a non-linear one, with many highly potent cells not necessarily having high doublet scores (**SI fig.4C**). Finally, we verified that similar results were obtained had we used another method for estimating doublet scores (**SI fig.S5**, **Methods**).

**Figure-4:**
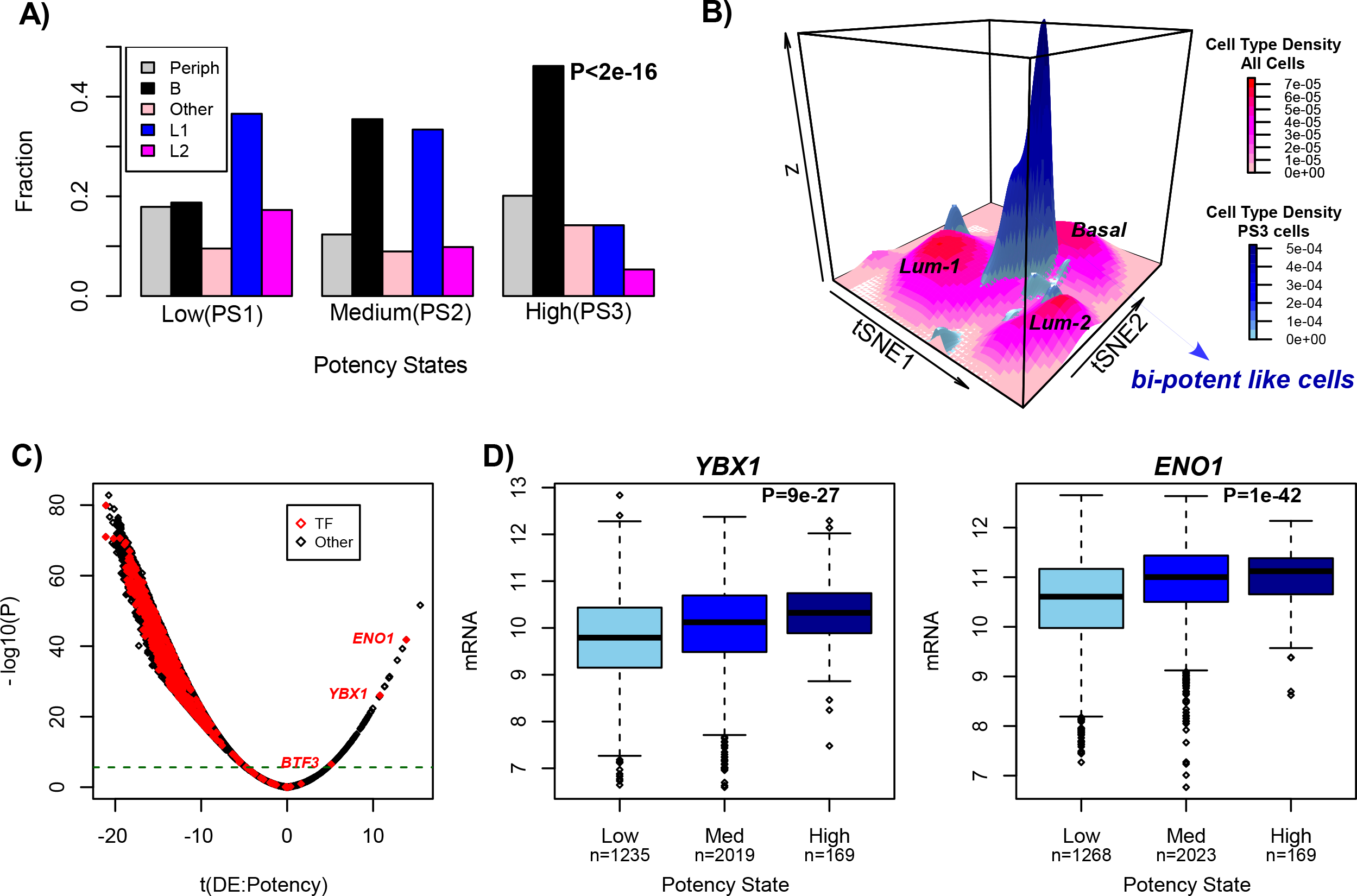
Integrated landscape reveals bi-potent state characterized by YBX1 and ENO1 expression. **A)** Percentage barplots displaying the relative distribution of breast epithelial subtypes (as inferred from the clustering using t-SNE + DBSCAN) among inferred potency states (Low, Medium, High). Single cells have been divided up into whether they clustered into the basal compartment (B), into the luminal-1 cluster (L1), the luminal-2 cluster (L2), all other clusters (Other) or whether they were not assigned into any cluster, defining peripheral cells (Periph). P-value is from a Kruskal-Wallis test to assess if the distribution of subtypes differs significantly within the high potency state. **B)**Surface cell-density map of all single cells (magenta colored surfaces) with the corresponding surface cell-density map of highly potent (PS3) cells superimposed (blueish colored surfaces). The x and y-coordinates label the t-SNE1 and t-SNE2 axes. The height of the surfaces (z) is a measure of cell-density in the x-y plane and is further indicated by different color tones. The z-axis is therefore not a measure of cell potency. **C)**Volcano plot of differential expression associated with potency, with x-axis labeling the t-statistic and y-axis labeling the statistical significance. Horizontal bar denotes the Bonferroni threshold, and red points indicate transcription factors (TFs). **D)**Boxplots of normalized log-expression (y-axis) for *YBX1* and *ENO1* against inferred potency state (x-axis) for all single cells where these genes were expressed. Numbers of single-cells assigned to each potency state is given. P-value is from a two-tailed linear regression. All single-cell cells derive from one individual (Ind-4).

### Bipotent cells are marked by *YBX1* and *ENO1* overexpression

In order to characterize the highly potent cells we performed DE analysis between high and low potent cells, irrespective of their epithelial subtype. The great majority of genes were downregulated in the more potent cells, with only 72 exhibiting overexpression (Bonferroni adjusted P<0.05, **Fig.4C**). Correspondingly, among the 1369 TFs, 582 exhibited differential expression (Bonferroni adjusted P < 0.05) with only 3 TFs (*ENO1, YBX1* and *BTF3*) exhibiting higher expression in the more potent cells (**Fig.4C-D**). Remarkably, *YBX1* and *ENO1* are two transcription factors whose targets are highly enriched for breast cancer GWAS eQTLs (Castro et al. 2016), thus implicating them in breast cancer risk. In addition, siRNA against *YBX1* in a normal ER- cell-line (MCF10A) resulted in significantly reduced cell-confluence and growth, even when compared to other breast cancer risk TFs (Castro et al. 2016). We confirmed that the associations of *YBX1* and *ENO1* expression with potency remained after adjustment for cell-cycle phase (**SI fig.S6, Methods**), and that their expression also correlated with cell potency in the scRNA-Seq data from the other 3 women (**SI fig.S7**).

### Upregulated bipotent single-cell signature correlates with mammary stemness

If the highly potent cells are bipotent, the expectation would be that they are transcriptionally similar to previously characterized mammary stem cells. We performed rank-based GSEA (Subramanian et al. 2005) on the 72 genes upregulated in the highly potent single cells to further characterize the putative bipotent cells. This revealed strong enrichment for ribosomal genes, but importantly also for genes upregulated in mammary stem-cells (**SI fig.S8**). In particular, we observed a relatively strong enrichment (12 gene overlap, OR=39, BH-adjusted Fisher-test P<1e-10) with a previously characterized mammary stem-cell signature (Pece et al. 2010). Of note, among the 12 overlapping genes, 9 (*RPS2, RPS7, RPS10, RPL8, RPS18, RPS3, RPL10A*) were ribosomal proteins or ubiquitin ribosomal fusion proteins (*UBA2 & FAU)*, consistent with recent findings that expression of ribosomal proteins is a universal marker of stemness and potency (Athanasiadis et al. 2017; Teschendorff and Enver 2017). Among the other 3 genes, we observed *NACA*, a protein that associates with the upregulated transcription factor *BTF3*, and *TXN* (thioredoxin), a protein involved in the response to intracellular nitric oxide.

To confirm the results of the GSEA, we obtained and normalized mRNA expression data from mammosphere-derived FACS sorted pools of quiescent mammary stem-cells and transit-amplifying progenitors (Pece et al. 2010) (**Methods**). Validating the association with stemness, the 12-genes exhibited increased expression in three separate pools of quiescent mammary stem-cells compared to their derived transit-amplifying progenitors (**Fig.5A-B**, Wilcox test P=0.001, **Methods**), a result which remained significant compared to randomly selected genes (**Fig.5C**, Monte Carlo P=0.0001). Results remained significant had we used all 72 genes (63 genes had representation on the Affymetrix platform used in Pece et al (Pece et al. 2010)) from the upregulated scRNA-Seq bipotent signature (**SI fig.S9**). However, interestingly, *YBX1* and *ENO1* were not upregulated in the quiescent mammary stem cells compared to the transit-amplifying progenitor cells (**SI fig.S9**), suggesting that while potency/stemness is marked by the expression level of ribosomal proteins, the progenitor non-quiescent state is associated with higher expression of *YBX1* and *ENO1*.

**Figure-5:**
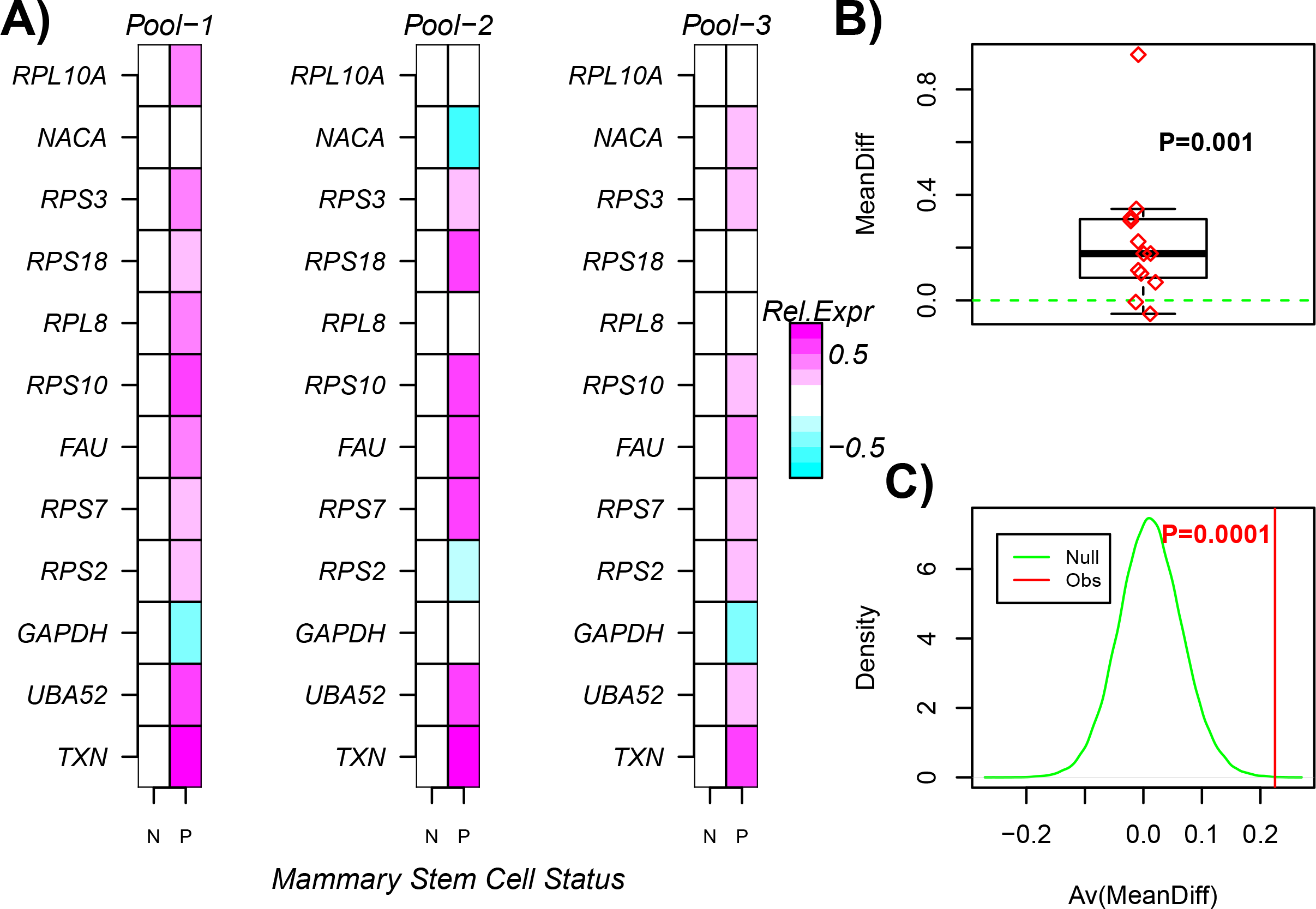
Bipotent single-cell expression signature is enriched for mammary stem cell genes. A) Normalized relative expression heatmaps for 12 represented genes from the 17-genes upregulated in the putative bipotent single-cells and which overlap with a mammary stem-cell signature, in 3 separate pools of FACS sorted quiescent mammary stem-cells (P) and their derived proliferative non-stem like progeny (N). **B)**Average expression difference between the P and N cells, averaged over the 3 separate pools. P-value is from a one-tailed Wilcoxon rank sum test. **C)**Monte-Carlo randomization analysis, where in each of 100,000 random selections of 17 genes, the average difference over the 3 pools is computed (green curve) and compared to the observed average difference (red, panel-B). Monte-Carlo P-value is given.

### *YBX1* expression correlates with luminal subtype and is increased in luminal progenitors

The correlation of *YBX1* expression with potency was particularly evident in the luminal compartment (**Fig.6A**, **SI fig.S10**), pointing towards *YBX1* as playing not only a key role in defining a basal progenitor phenotype, but also potentially as a luminal progenitor. We were able to further validate this in two ways. First, its expression was also higher in the more immature luminal alveolar-like phenotype, in line with the fact that these alveolar luminal cells should be more enriched for progenitors (**Fig.6B**). Second, using bulk expression data from FACS sorted luminal progenitor and differentiated luminal cells (Shehata et al. 2012), we found *YBX1* expression to be highest for the EpCAM+/CD49f+/ALDH+ population (**Fig.6C**, **Wilcox test P=0.003**), which defines the most likely luminal progenitor phenotype, or at least the one that gives rise to milk-producing alveolar cells (Shehata et al. 2012).

**Figure-6:**
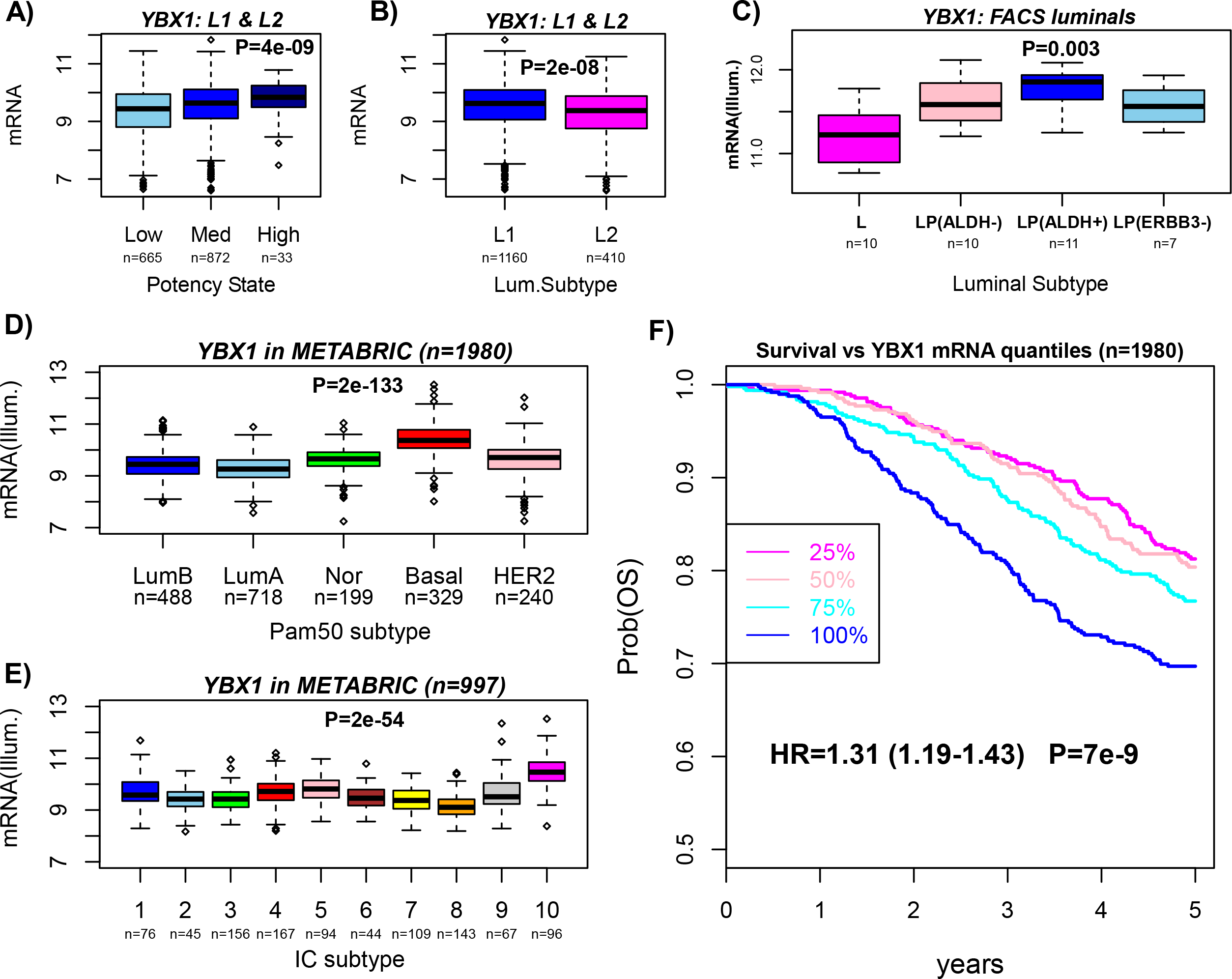
YBX1 expression characterizes luminal progenitors and basal breast cancer. **A)** Boxplots of normalized log-expression (y-axis) for *YBX1* against inferred potency state (x-axis) for all single cells assigned to luminal L1 and L2 clusters and where *YBX1* is expressed. Numbers of single-cells in each group is given. P-value is from a linear regression. **B)** Boxplots of normalized log-expression (y-axis) for *YBX1* against luminal cluster, using only single cells where *YBX1* is expressed. Numbers of single-cells in each group is given. P-value is from a one-tailed Wilcox test. **C)**Boxplots of Illumina normalized log-expression (y-axis) for *YBX1* against luminal subtype as defined by FACS-sorting (x-axis): L=differentiated non-clonogenic luminal, LP(ALDH-)=ALDH- luminal progenitor, LP(ALDH+)=ALDH+ luminal progenitor. LP(ERBB3-)=ERBB3- and ALDH- luminal progenitor. P-value is from a one tailed Wilcox-test comparing LP(ALDH+) to all others. **D)**Boxplots of normalized Illumina log-expression (y-axis) for *YBX1* against PAM50 intrinsic subtype in the full METABRIC cohort. P-value is from a Kruskal-Wallis test**. E)** As D), but for the integrative cluster (IC) subtypes (available in discovery set only). **F)**Kaplan Meier overall survival curves for *YBX1* expression, stratified by quantiles of *YBX1* expression, and censored at 5 years after diagnosis. Hazard Ratio (HR), 95% CI and P-value are from a Cox proportional hazards regression.

Of note, we obtained similar results if instead of *YBX1* we used the earlier 17-gene or 72-gene signatures marking the bipotent cells. Indeed, the great majority of these genes were observed to be overexpressed in the EpCAM+/CD49f+/ALDH+ population compared to all other cell populations, a result which was highly significant as assessed using 100,000 Monte-Carlo randomizations (P<1e-5, **SI fig.S11**).

### *YBX1* expression marks basal breast cancer

Given that *YBX1* exhibited highest expression in the more potent single-cells, and that these were enriched within the basal compartment, it is natural to posit that *YBX1* may mark the cell of origin for basal breast cancer. If so, *YBX1* expression should be highest in basal breast cancer compared to other breast cancer subtypes. We were able to confirm this with high statistical significance within the METABRIC study (Curtis et al. 2012), which profiled almost 2000 primary breast cancers (**Fig.6D**). Similar results were obtained if instead of *YBX1* we used the complete 17-gene or 72-gene signatures marking the bipotent cells (**SI fig.S12**). In terms of the integrative cluster (IC) subtypes, as defined by METABRIC, *YBX1* expression was highest in IC-5 and IC-10 (**Fig.6E**). These two integrative cluster subtypes exhibited the worst disease-specific 5-year survival rates among all IC subtypes (Curtis et al. 2012). In line with this, we observed that *YBX1* expression also correlated with a poor clinical outcome (**HR=1.31, P=7e-9, Fig.6F**). However, the association with outcome was mainly driven by ER-status, since in an analysis stratified by ER-status we did not observe any significant association (HR=1.11, P=0.12 in ER+; HR=0.96, P=0.64 in ER-).

## Discussion

Here we have demonstrated “proof-of-concept” that our signaling entropy rate measure can be used to identify rare subpopulations of highly-potent cells, which may represent novel candidate progenitor or stem-like cells. Indeed, application to almost 4,000 single cells from the mammary epithelium identified a rare (5%) subpopulation of relatively high potency, which is likely to represent a mammary progenitor-like state. We extensively validated the potency assignments of the single-cells, and consistent with the prevailing view that most mammary progenitors are basal cells, the highly potent cells were over-represented within the basal compartment. The ability to stratify single cells into different potency states allowed us to infer and compare the cell-density surface maps for all potency states, revealing that highly potent cells exhibited a strikingly different landscape to those of lower potency, with the region of maximum cellular density defining a distinctive bi-modal ridge between the basal and alveolar luminal clusters, with the largest peak occurring within the basal compartment. Thus, without the need for any prior assumptions, LandSCENT predicts that these highly potent cells may represent a bi-potent subpopulation that gives rise not only to basal cells but also to luminal progenitors. Supporting this view, we found that the main TF characterizing these highly potent cells (*YBX1*) plays a key role in maintaining the self-renewal and proliferative capacity of basal cells (Castro et al. 2016) and that it is also overexpressed in FACS sorted luminal progenitor populations compared to luminal differentiated cells. In addition, we found that among the top-ranked genes upregulated in these putative bipotent cells, there was a clear and significant enrichment for genes that have been found to mark quiescent mammary stem cells and stemness generally (Athanasiadis et al. 2017; Teschendorff and Enver 2017). We stress that these independent validations using orthogonal expression data from bulk samples clearly shows that our results are not technical artefacts of single-cell data.

The significance of *YBX1* extends to the cancer-risk context. First, there is already substantial evidence demonstrating that *YBX1* transforms mammary epithelial cells, via binding to the *BMI1* promoter and chromatin remodeling, leading to basal breast cancer (Davies et al. 2014). In line with this, *YBX1* is also more highly expressed in basal breast cancer compared to all other breast cancer subtypes, consistent with it marking cells that give rise to basal breast cancer. Second, *YBX1* expression also marks luminal progenitor cells, and a subset of basal breast cancers, notably BRCA1 mutant ones, are thought to arise from a mis-programmed luminal progenitor (Lim et al. 2009; Shehata et al. 2012). Indeed, the single-cell landscape inferred with LandSCENT underscores the similarity of the highly potent cells within the basal compartment with those in the immature luminal cluster, strongly suggesting that the cell of origin for basal breast cancer may well be a bi-potent like cell that shares an expression profile similar to that of luminal progenitors, including notably *YBX1.* Third, *YBX1* has been shown to interact with *ESR1*, and via *FGFR2* signaling may contribute to tamoxifen resistance (Campbell et al. 2018). Fourth, it has been observed that genes within the *YBX1* regulon are strongly enriched for GWAS breast cancer eQTLs (Castro et al. 2016). This is a highly significant observation, given the growing evidence that molecular alterations (both inherited and somatic) affecting the adult stem/progenitor cells within the tissue is a main risk factor for epithelial cancer development (Tomasetti and Vogelstein 2015b; Tomasetti and Vogelstein 2015a; Yang et al. 2016; Zhu et al. 2016; Tomasetti et al. 2017). Thus, we speculate that it is the genetic and epigenetic alterations that accumulate within the bi-potent progenitor cell pool identified here, which may confer the risk of breast cancer, especially basal breast cancer.

In future, it will be important to conduct more comprehensive and deeper sequencing of single cells in the mammary epithelium in order to construct accurate expression profiles for the bi-potent cell pool identified here. In this regard, we point out that we were here severely limited by the relatively low coverage of the 10X Chromium data (an average of only ~60,000 reads per cell), which did not allow us to fully determine the differential expression landscape of the bi-potent cells. The identification of *YBX1* (and *ENO1*) is a promising start, but we anticipate that other regulators will also play a key role in defining these bi-potent cells. We envisage that the computational framework presented here will play an important role as a means of identifying and characterizing the bi-potent cells in the larger and deeper scRNA-Seq studies to be performed in the near future. Importantly, LandSCENT will be equally applicable to future large-scale scRNA-Seq studies performed on cancer tissue which aim to identify putative cancer-stem-cells (Tirosh et al. 2016a; Tirosh et al. 2016b; Teschendorff and Enver 2017).

In summary, we have presented a novel 3-dimensional clustering algorithm for scRNA-Seq data, which uses an unbiased and assumption-free approach to estimate cell potency, and which is used to perform single-cell clustering within each potency state. Application of this simple yet powerful approach to scRNA-Seq data from the mammary epithelium naturally predicts a bipotent cluster, which as shown here is characterized by regulators that have been shown to modulate breast cancer risk. This study therefore provides a link between the progenitor and stem like cell population that controls homeostasis within a complex epithelial tissue and regulatory factors implicated in cancer risk of that same tissue. Our algorithm and findings may serve as a general paradigm for analogous studies in other tissue types.

## Methods

### Single cell data and preprocessing

The scRNA-Seq data analysed in this work derives from the study of Nguyen et al (Nguyen et al. 2018), who used the 10X Genomics Chromium platform to sequence a total of 24,646 cells from reduction mammoplastic specimens from 4 separate nulliparous women (Ind4-7), at an average read-depth of 60,000 reads per cell. Mapped read count data from the 4 individuals was downloaded from GEO (GSE113197), and further normalized as follows: for each cell we counted the number of expressed genes (“coverage per cell”), and for each gene we also counted the number of times it was expressed across all single cells (“coverage per gene”). For each cell, we also computed the total read count mapping to mitochondrial genes, which revealed low cell coverage for those cells having a high proportion of mitochondrial gene read counts. Based on this, we selected all cells expressing at least 1000 genes and with the proportion of mitochondrial read counts less than 0.05, leaving a total of 23,369 cells. Mitochondrial genes were removed and the total read count per cell *c* recomputed (*TRC*_*c*_). Denoting the maximum of *TRC*_*c*_ by *maxC*, and the read count matrix by *RCM*, the latter was normalized with the following transformation: *LSC*_*gc*_=*log*_*2*_*(RCM*_*gc*_\**maxC*/*TRC*_*c*_ + *1.1)*. Finally, we only use Entrez gene ID annotated genes, which resulted in a log-normalized single cells matrix of dimension 22049 genes and 23369 cells (3473 for Ind-4, 6811 for Ind-5, 5807 for Ind-6 and 7278 for Ind-7).

### The <u>Land</u>scape <u>S</u>ingle-<u>C</u>ell <u>En</u>tropy and Cell-<u>T</u>ype (LandSCENT) algorithm

LandSCENT is a direct extension of the SCENT algorithm. There are three steps to the LandSCENT algorithm: (1) Inference of potency states: estimation of the differentiation potency of single cells via computation of the signaling entropy rate (SR) and subsequent inference of the potency state distribution across the single cell population. (2) Inference of cell-types: we perform t-SNE (van der Maaten 2008) followed by density-based spatial clustering (dbscan) (Ester et al. 1996) on a suitably dimensionally reduced *LSC* matrix. (3) Construction of an integrated landscape defined over potency-states and cell-types using cell-density surface maps to reveal cellular-states. We note that step-1 is the exact same procedure as used in our original SCENT algorithm (Teschendorff and Enver 2017).

#### Step-1 Inference of potency states

We estimate differentiation potency of each single cell by computing the signaling entropy using the same prescription as used in our previous publications (Banerji et al. 2013; Teschendorff et al. 2014). Briefly, the normalized genome-wide gene expression profile of a sample (this can be a single cell or a bulk sample) is used to assign weights to the edges of a highly curated protein-protein interaction (PPI) network. The construction of the PPI network itself is described in detail elsewhere (Banerji et al. 2013), and is obtained by integrating various interaction databases which form part of Pathway Commons (www.pathwaycommons.org) (Cerami et al. 2011). The weighting of the network via the transcriptomic profile of the cell provides the biological context. The weight of an edge between protein *i* and protein *j*, denoted by *w*_*ij*_, is assumed to be proportional to the normalized expression levels of the coding genes in the cell, i.e. we assume that *w*_*ij*_ ~ *x*_*i*_*x*_*j*_. We interpret these weights (if normalized) as interaction probabilities. The above construction of the weights is based on the assumption that in a sample with high expression of *i* and *j*, that the two proteins are more likely to interact than in a sample with low expression of *i* and/or *j*. Viewing the edges generally as signaling interactions, we can thus define a random walk on the network, assuming we normalize the weights so that the sum of outgoing weights of a given node *i* is 1. This results in a stochastic matrix, *P*, over the network, with entries

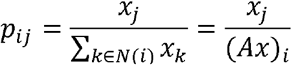

where *N(i)* denotes the neighbors of protein *i*, and where *A* is the adjacency matrix of the PPI network (*A*_*ij*_=*1* if *i* and *j* are connected, 0 otherwise, and with *A*_*ii*_=0). The signaling entropy is then defined as the entropy rate (denoted *Sr*) over the weighted network, i.e.

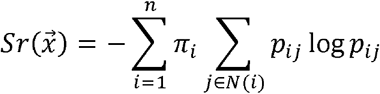

where *π* is the invariant measure, satisfying *πP*=*π* and the normalization constraint *π*^*T*^**1**=1. The invariant measure, also known as steady-state probability, represents the relative probability of finding the random walker at a given node in the network (under steady state conditions i.e. long after the walk is initiated). Nodes with high values thus represent nodes that are particularly influential in distributing signaling flux in the network. In the steady-state we can assume detailed balance (conservation of signaling flux, i.e. *π*_*i*_*p*_*ij*_ = *π*_*j*_*p*_*ji*_), and it can be shown (Teschendorff et al. 2014) that *π*_*i*_ = *x*_*i*_(*Ax*)_*i*_/(*x*^*T*^*Ax*). Given a fixed adjacency matrix *A* (i.e. fixing the topology), it can also be shown (Teschendorff et al. 2014) that the maximum possible *Sr* among all compatible stochastic matrices *P*, is the one with 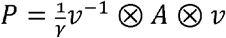 where ⊗ denotes product of matrix entries and where *v* is the dominant eigenvector of *A*, i.e. *Av*=λ*v* with λ the largest eigenvalue of *A*. We denote this maximum entropy rate by *maxSr*, and define the normalized entropy rate (with range of values between 0 and 1) as

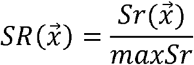

Since *SR* is bounded between 0 and 1, we next transform the *SR* value of each single cell into their logit-scale value, i.e. *y(SR)*=*log*_*2*_*(SR/(1-SR))*. Subsequently, we fit a mixture of Gaussians to the *y(SR)* values of the whole cell population, and use the Bayesian Information Criterion (BIC) (as implemented in the *mclust* R-package) (Yeung et al. 2001) to estimate the optimal number *K* of potency states, as well as the state-membership probabilities of each individual cell. Thus, for each single cell, this results in its assignment to a specific potency state.

#### Step-2 Inference of cell-types

Cell-types are inferred as significant clusters using cell-density in the two-dimensional t-SNE space as the main criterion. Preliminary dimensional reduction is achieved by first selecting genes with a mean average expression larger than 1, and also a standard deviation larger than 1. These thresholds were chosen after inspection of the mean-variance plot, and in the case of Ind-4 this resulted in 4261 highly variable and expressed genes. To map the high dimensional nature of the data matrix to a two-dimensional subspace we used t-SNE with an initial dimension of 30, a perplexity parameter of 30, 1000 maximum iterations and epoch parameter set to 100. We then used the dbscan algorithm (density-based spatial clustering) with eps=5 and minPts=15 to identify significant clusters. Thus, after steps-1 and 2, each cell is assigned to a unique potency state and co-expression cluster (cell-type).

#### Step-3 Inference and construction of an integrated landscape of cell-states

Finally, we construct cell-density surface maps for all single cells within each of the inferred potency states. In these surface maps, the elevation is directly proportional to cell-density. By comparing the resulting landscapes for each potency state, this may reveal novel cellular states, defined by both potency and expression subtype.

### Estimation of cell-cycle and TPSC pluripotency scores

To identify single cells in either the G1-S or G2-M phases of the cell-cycle we followed the procedure described in (Tirosh et al. 2016a). Briefly, genes whose expression is reflective of G1-S or G2-M phase were obtained from (Whitfield et al. 2002; Macosko et al. 2015). A given normalized scRNA-Seq data matrix for a given individual is then z-score normalized for all genes present in these signatures. Finally, a cycling score for each phase and each cell is obtained as the average z-scores over all genes present in each signature. When adjusting differential expression analyses for cell-cycle phase, we included the G1-S and G2-M scores as covariates in the linear models.

### Bulk expression datasets

In this study we used three mRNA expression datasets from bulk samples. One dataset consists of 38 FACS sorted bulk samples (Illumina expression beadarrays), as profiled by Shehata et al (Shehata et al. 2012). Of the 38 samples, 10 were categorized as luminal non-clonogenic (L), i.e. terminally differentiated cells, with the rest (n=28) making up a relatively differentiated (EpCAM+/CD49f+/ALDH-, n=17) and undifferentiated (EpCAM+/CD49f+/ALDH+, n=11) luminal progenitor (LP) populations. The two undifferentiated LP populations were further distinguished by expression or not of ERBB3. mRNA expression data was generated using Illumina Beadarrays and we used the normalized data, as described in (Shehata et al. 2012).

The second dataset is the METABRIC study, which profiled almost 2000 primary breast cancers using Illumina expression beadarrays (Curtis et al. 2012). We used the assignment of tumors to PAM50 intrinsic and integrative cluster (IC) subtypes as given by the METABRIC study. We used the normalized data, as provided by the METABRIC consortium.

A third Affymetrix mRNA expression dataset derives from Pece et al (Pece et al. 2010). This set consists of 3 separate pools of FACS sorted cell populations. Each pool contains a quiescent putative mammary stem cell population, as well as a population of derived progeny, consisting of transit-amplifying progenitor cells, thus a total of 6 bulk samples. We normalized the HGU133 plus2 data using the affy BioC package, specifically, the rma function. Only probes mapping to an Entrez gene ID were used, data was quantile normalized using limma, and probes mapping to the same gene were averaged, resulting in a normalized data matrix over 20186 genes and 6 samples.

### Differential Expression Analysis

When performing differential expression analysis on single-cell data, for each gene we always restrict to those cells where the gene is expressed. That is, we remove all dropouts and don’t impute data. When correlating to potency, we used a linear model between the normalized expression profile and the potency estimates, optionally adjusting for the two cell-cycle scores computed earlier. In the case of the Illumina beadarray datasets, we used the normalized data from the respective publications (Curtis et al. 2012; Shehata et al. 2012) and called DE using the empirical Bayes limma framework (Smyth 2004). We always use Bonferroni-adjusted thresholds to call statistical significance unless there are too few hits, in which case we relax the threshold using FDR<0.05 instead.

### Doublet score analysis

We used two different simulation-based methods to derive doublet scores for each cell and to identify those more likely to be doublets. One approach used the simulation method of Dahlin et al (Dahlin et al. 2018) to obtain doublet scores for all single cells that passed QC and for each individual separately. Specifically, we used the doubletCells function (using approximate=TRUE option) from the *scran* R-package (version 1.10.1) (Lun et al. 2016). In the second approach we used the Python package Scrublet (Wolock et al. 2018) (doi: https://doi.org/10.1101/357368). Within Scrublet, the scrub_doublets function, which is responsible for computing doublet scores and predicting doublets within a dataset, was run using default parameters.

### Code Availability

SCENT is freely available as an R-package from github: https://github.com/aet21/SCENT

### Data Access

Data analyzed in this manuscript is already publicly available from the following GEO (www.ncbi.nlm.nih.gov/geo/) accession numbers: GSE113197, GSE35399, GSE18931 or from the EGA (http://www.ebi.ac.uk/ega/) accession number EGAS00000000083.

## Supporting information

## Acknowledgements

The authors would like to thank Devon Lawson for useful discussions and Carlos Caldas for provision of METABRIC data. This work was supported by NSFC (National Science Foundation of China) grants, grant numbers 31571359 and 31401120 and by a Royal Society Newton Advanced Fellowship (NAF project number: 522438, NAF award number: 164914).

