## Supplementary material for "Integrated single-cell potency and expression landscape in mammary epithelium reveals novel bipotent-like cells associated with breast cancer risk"

### SUPPLEMENTARY FIGURES

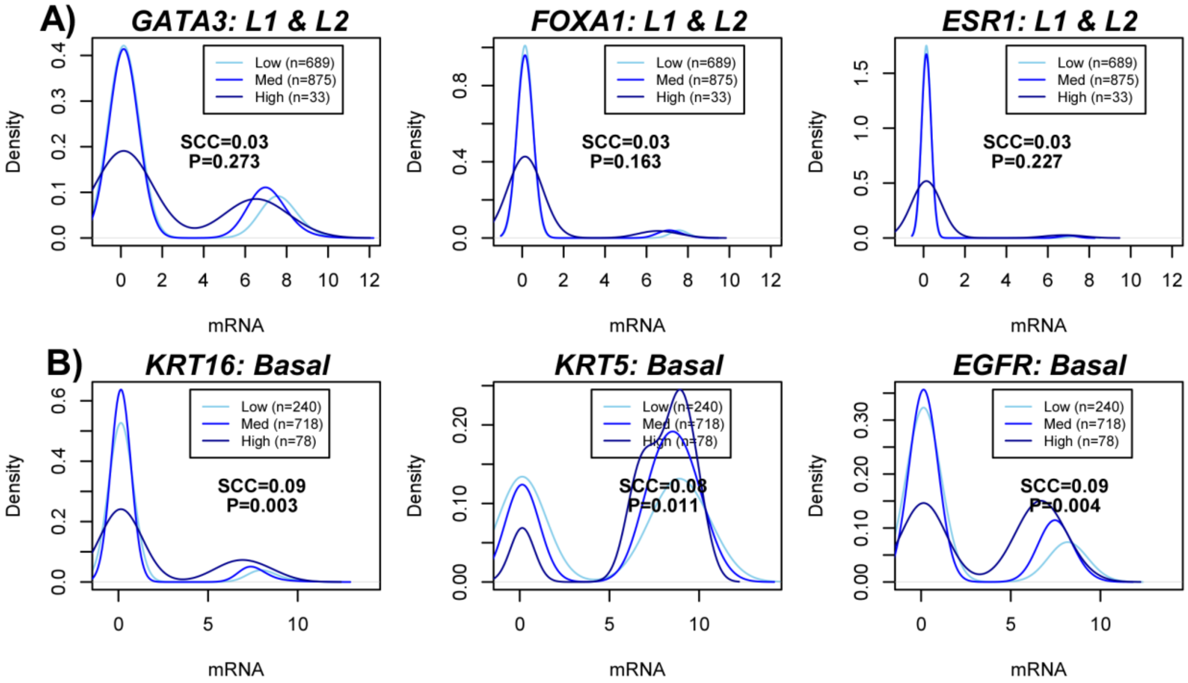


**fig.S1: Density profiles of luminal and basal marker expression. A)** For 3 well-known luminal differentiation markers we contrast the density distributions of their expression levels across the three inferred potency states, as indicated. SCC denotes the Spearman rank correlation coefficient between expression and potency and P is the corresponding two-tailed P-value. **B)** As A), but now for 3 well-known basal markers.

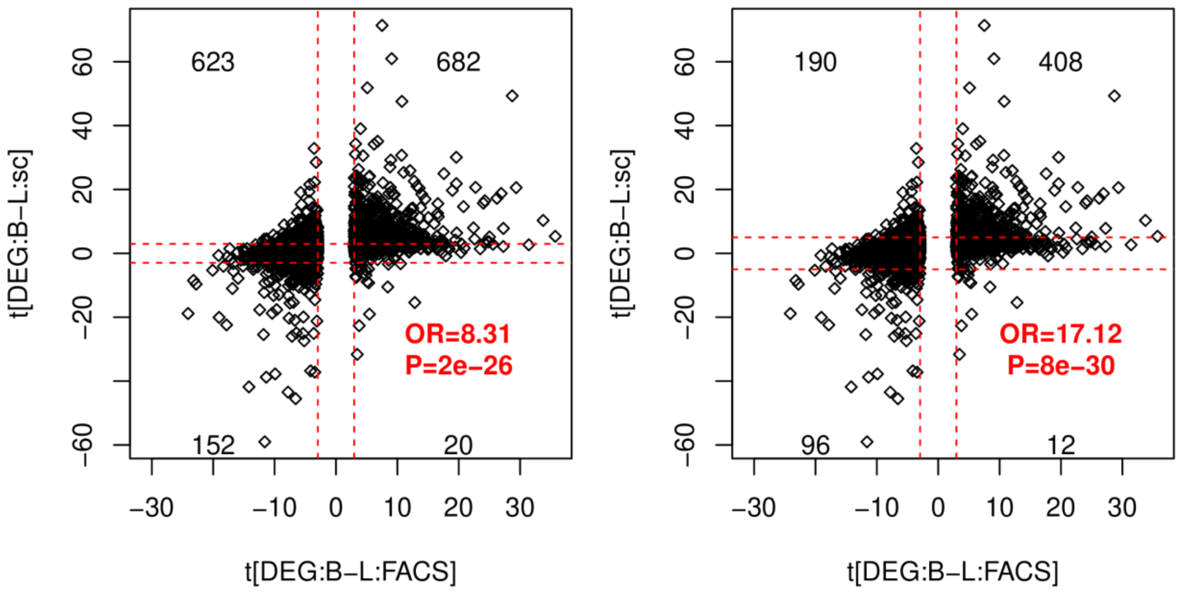


**fig.S2: Validation of DEG calling method using independent FACS bulk data.** Each panel depicts a scatterplot of the t-statistics of differential expression between FACS sorted basal and luminal cell populations (Illumina beadarray data) (x-axis) against the corresponding t-statistics of differential expression between the single-cell basal and luminal (L1&L2) clusters (y-axis). For the left panel, the red dashed vertical lines indicate the FDR=0.05 threshold, whereas the horizontal lines indicate the corresponding t-tstatistic threshold. For the right panel, the horizontal lines indicate a more stringent threshold. Odds Ratios and one-tailed Fisher-exact test P-values are given.

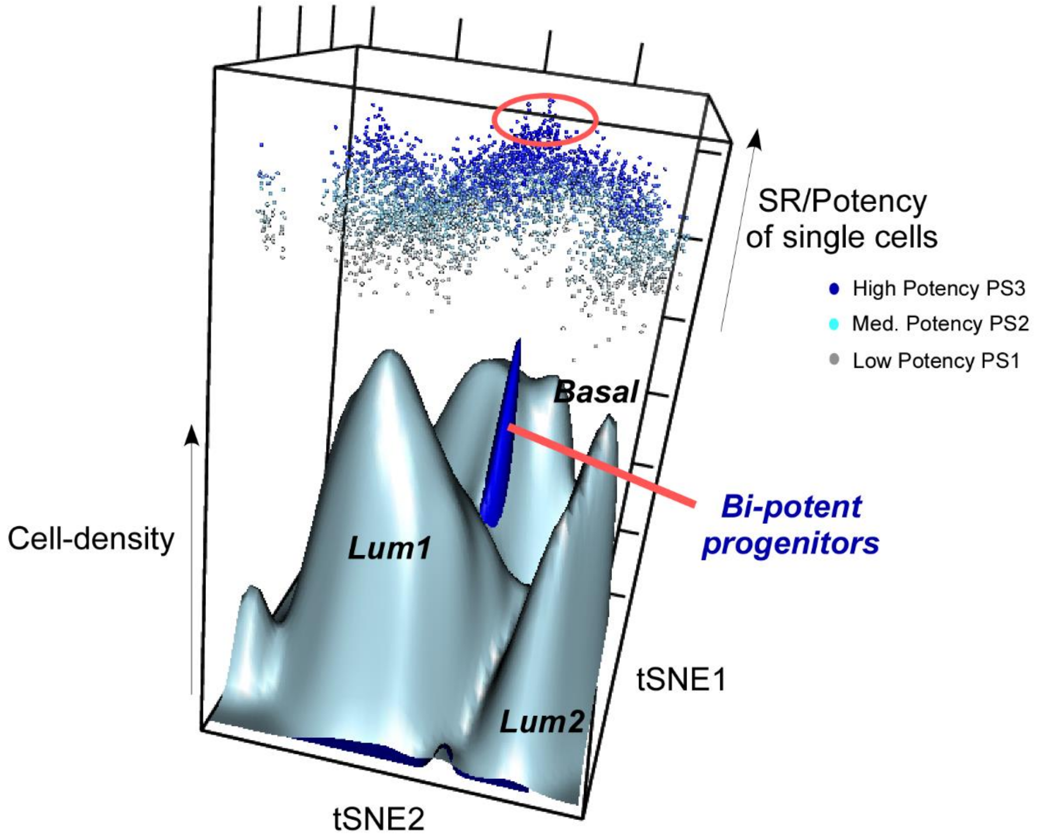


**fig.S3: Cell density surface map with single-cell entropies superimposed.** Three-dimensional representation of cellular density of the 3473 single cells, with x-y plane defined by the t-SNE coordinates and with z-axis labeling the cell-density (grey surface maps). Superimposed is the cell-density surface map for the cells categorized into high potency (blue surface map), defining a peak in between the basal and luminal-1 clusters and defining putative bi-potent progenitor. Also superimposed are the single-cell entropy/potency values (SR) for all 3473 single cells, colored by their inferred potency class and with their elevation directly proportional to SR.

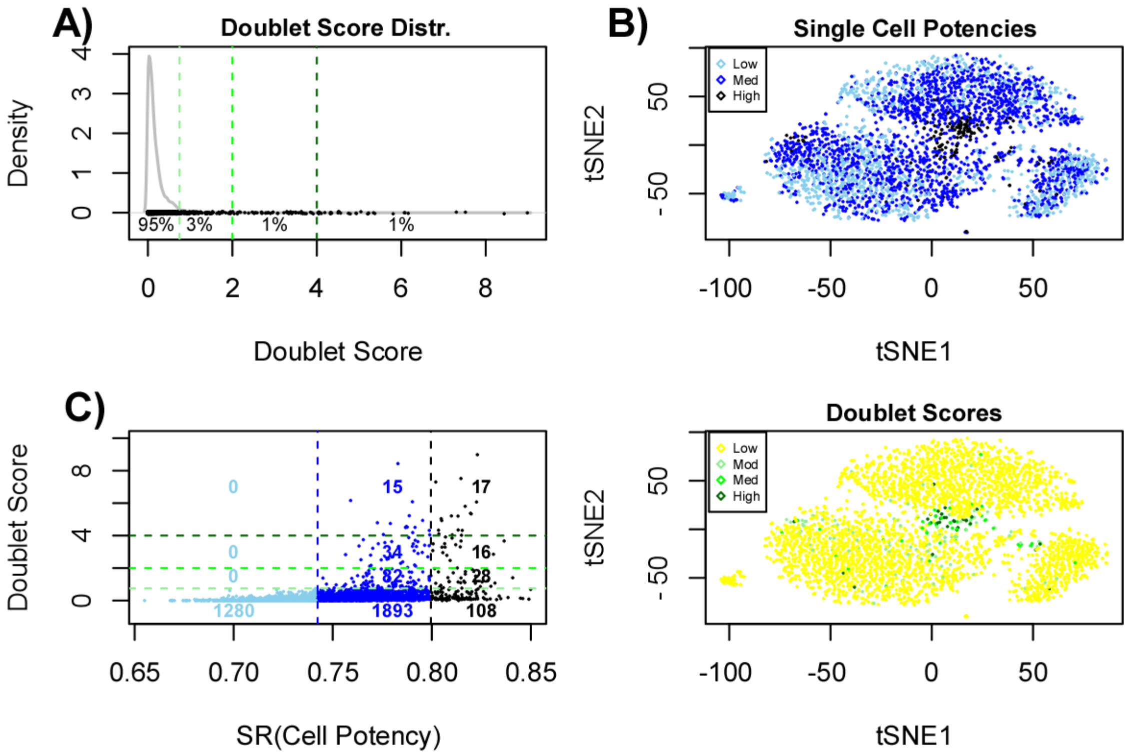


**fig.S4: Doublet analysis using scran. A)** Density distribution of doublet scores of the 3473 single cells from individual-4, with the proportions of cells falling within different score-bin categories. Doublet scores were obtained using the *scran* R-package. **B)** tSNE plot for all 3473 single cells, displaying their inferred potency states (upper panel) and their binned doublet scores (lower panel), as shown. **C)** Scatterplot of the doublet scores (y-axis) against signaling entropy rate (SR, x-axis) for all 3473 single cells. Dashed lines indicate the boundaries defining potency states and binned doublet scores, and the numbers indicate the number of cells falling within each rectangular bin.

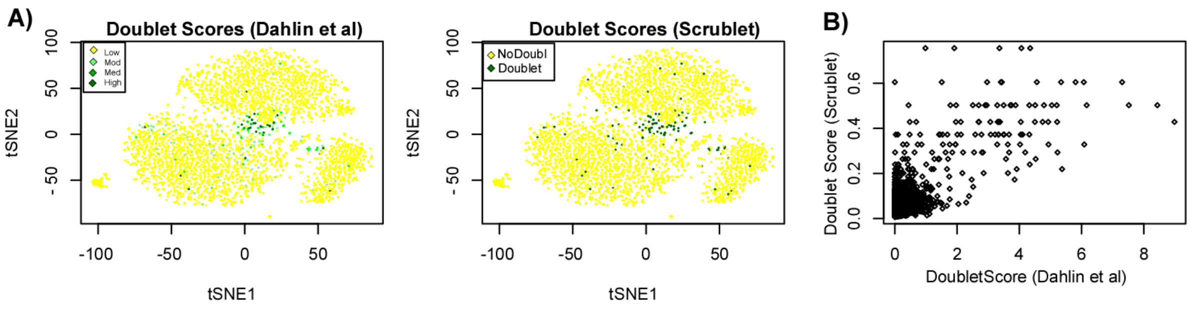


**fig.S5: Doublet analysis using scrublet. A)** tSNE plots for all 3473 single cells from individual-4, displaying their doublet scores according to scran (left panel) and scrublet (right panel). **B)** Scatterplot of the doublet scores according to scran (x-axis) against those of scrublet (y-axis) for all 3473 single cells.

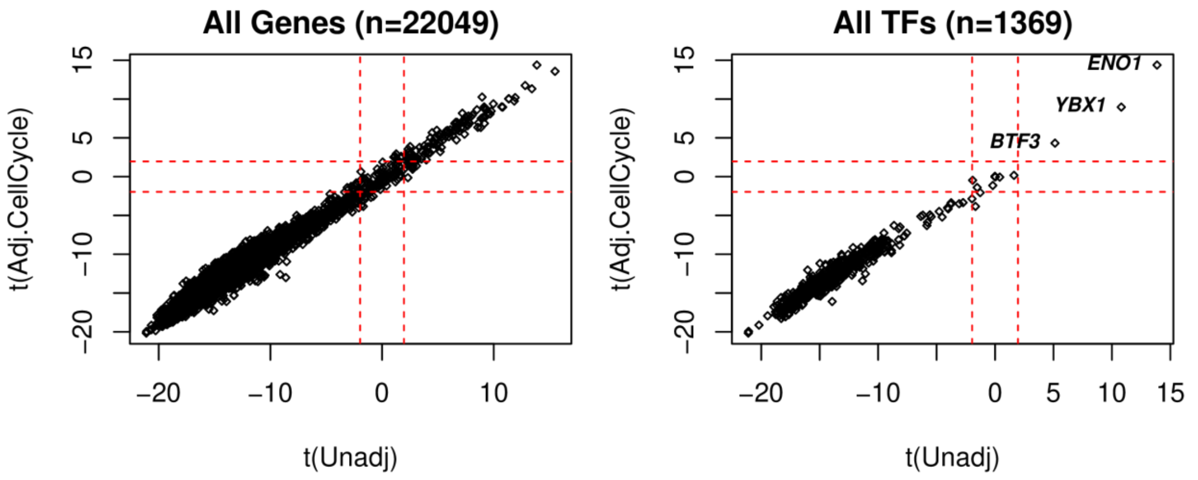


**fig.S6: Differential expression potency analysis adjusted for cell-cycle phase.** Scatterplots of t-statistics of association with signaling entropy rate and gene expression unadjusted for cell-cycle phase (x-axis) vs adjusted for cell-cycle phase (y-axis) for all genes (left panel) and for all transcription factors (TFs) (right panel).

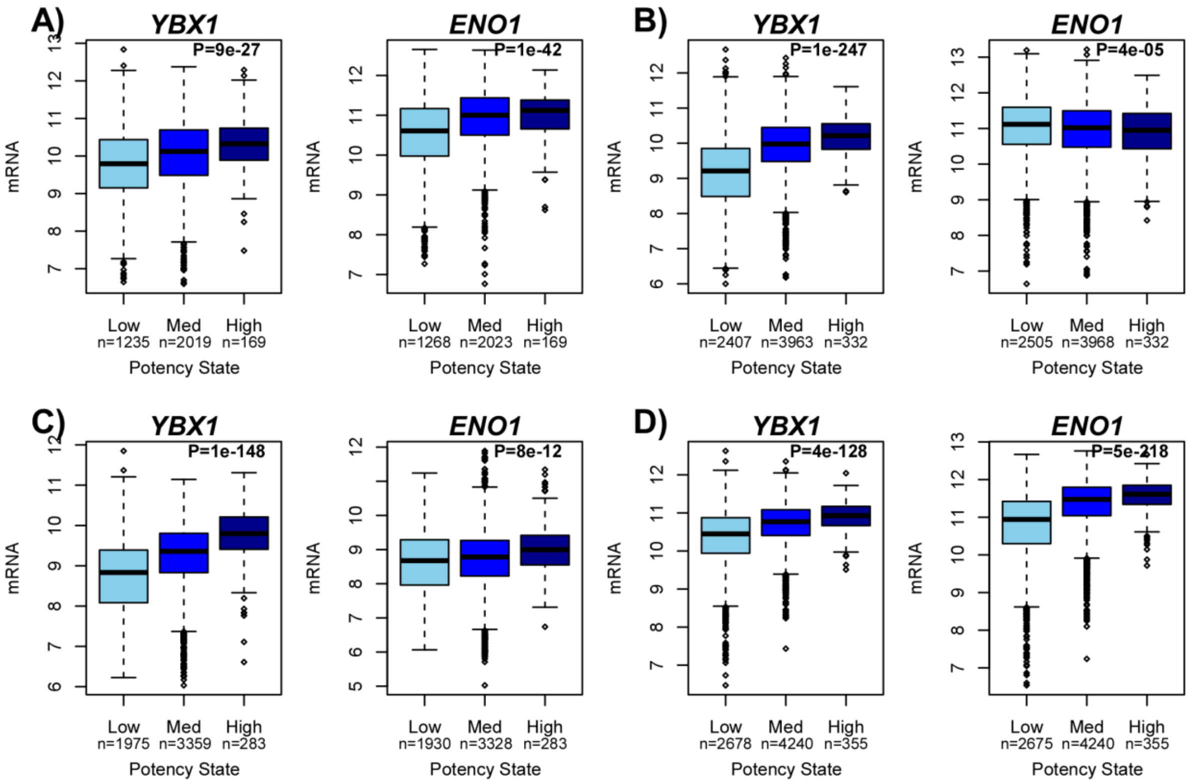


**fig.S7: YBX1 and ENO1 expression correlate with potency. A)** Boxplots of normalized log-expression (y-axis) for *YBX1* and *ENO1* against inferred potency state (x-axis) for all single cells where these genes were expressed. Numbers of single-cells assigned to each potency state is given. P-value is from a two-tailed linear regression. All single-cell cells derive from one individual (Ind-4). **B, C, D)** As A), but for three different individuals.

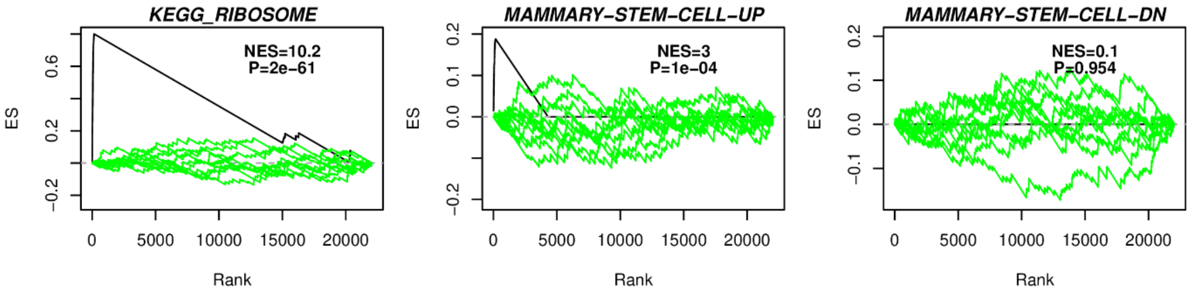


**fig.S8: Rank-based GSEA reveals correlation of mammary stem-cell and ribosomal protein modules with increased potency in mammary epithelium.** Plots of the Enrichment Score (ES, y-axis) from rank-based GSEA against rank index position (x-axis) for genes ranked according to their positive correlation with potency as assessed using the scRNA-Seq data (black line), and for three different biological terms from the MSigDB dataset: Ribosomal genes from the KEGG database, genes upregulated in mammary stem cells (Pece et al) and genes downregulated in mammary stem cells (Pece et al). Green curves describe dependence of the ES score on rank position after Monte-Carlo randomization of the gene-ranking, for 10 different Monte-Carlo runs. The Normalized Enrichment Score (NES) defined by the ratio of the observed maximum ES score to the mean of the maximum over 1000 Monte-Carlo runs is given, as well as the associated P-value derived by approximating the max ES scores over the 1000 Monte-Carlo runs as a Gaussian.

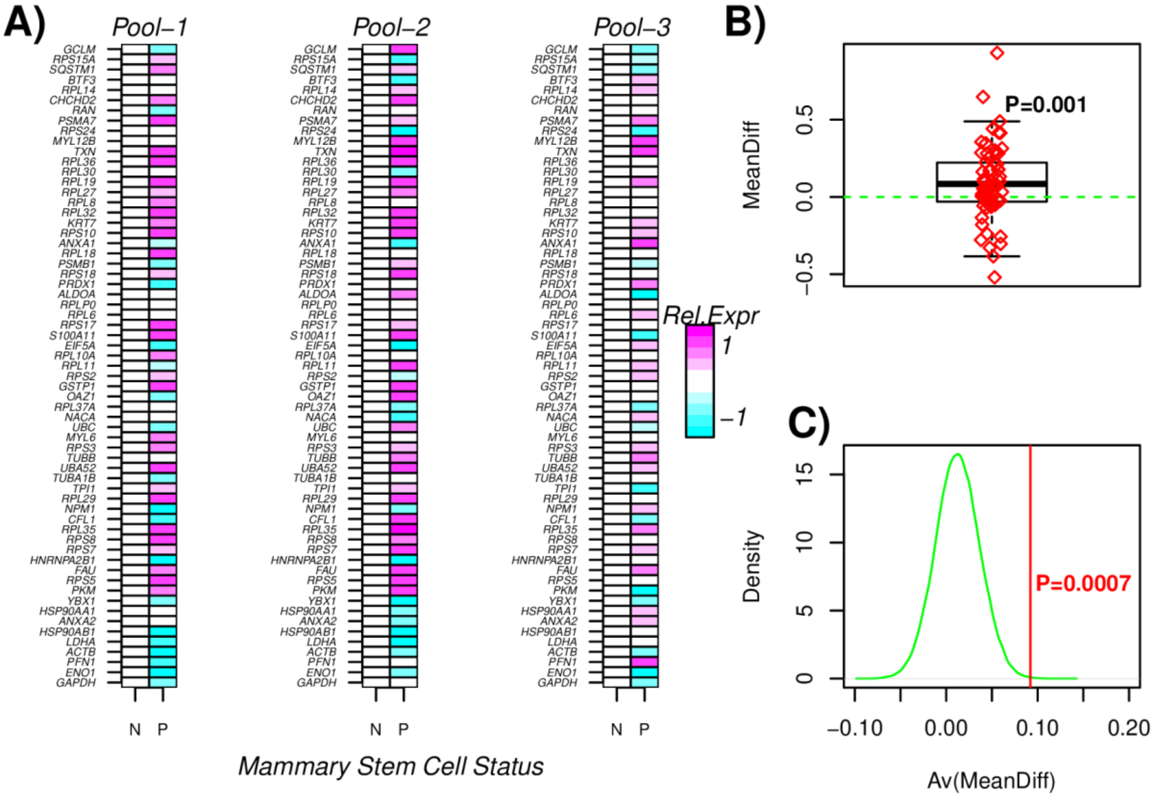


**fig.S9: Bipotent single-cell expression signature is increased in mammary stem cell pools.** **A)** Normalized relative expression heatmaps for 63 represented genes from the 72 genes upregulated in the putative bipotent single-cells, in 3 separate pools of FACS sorted quiescent mammary stem-cells (P) and their derived proliferative non-stem like progeny (N). **B)** Average expression difference between the P and N cells, averaged over the 3 separate pools. P-value is from a one-tailed Wilcoxon rank sum test. **C)** Monte-Carlo randomization analysis, where in each of 100,000 random selections of 63 genes, the average difference over the 3 pools is computed (green curve) and compared to the observed average difference (red line). Monte-Carlo P-value is given.

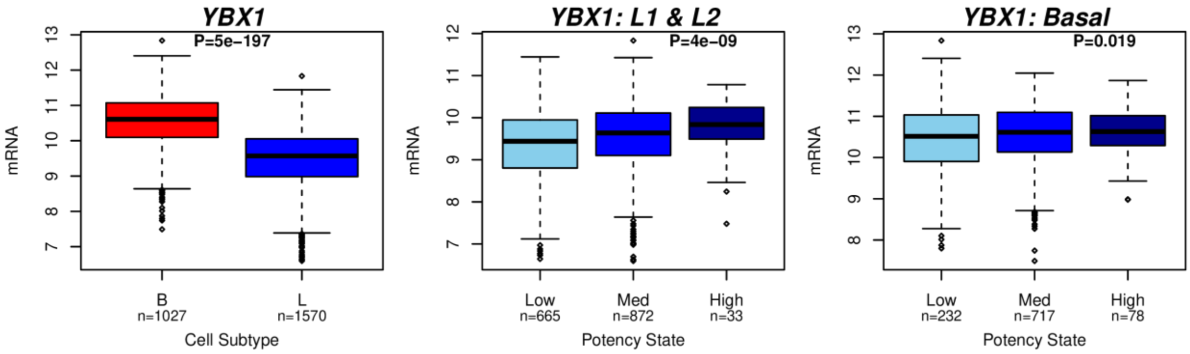


**fig.S10: YBX1 expression across the basal luminal divide and potency states. Left panel:** Boxplot of expression of YBX1 between the single-cells from the basal cluster (B) and those of the two luminal (L1 & L2) clusters. P-value is from a t-test and number of cells in each category is given below boxplot. **Middle & right panels:** Boxplot of YBX1 expression against potency state restricting to luminal and basal cells, respectively. P-value is from a linear regression t-test and number of cells in each category is given below boxplot. In all panels, only cells expressing *YBX1* were included.

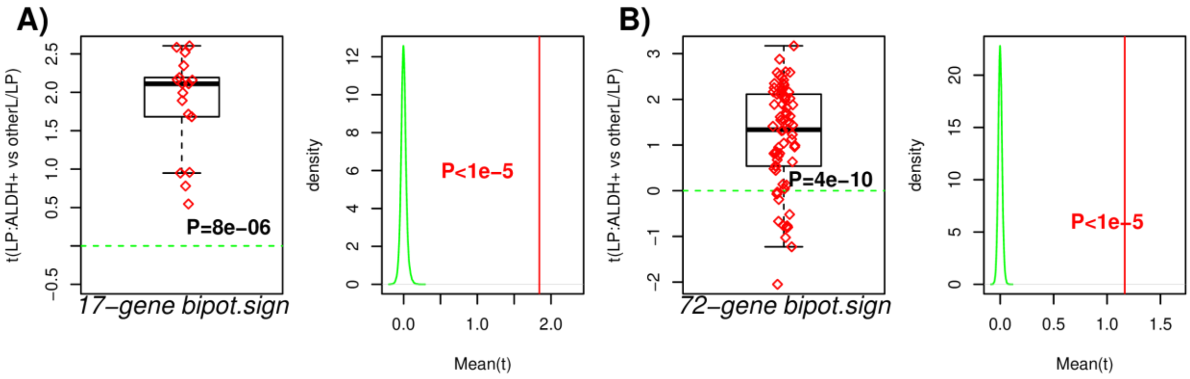


**fig.S11: Correlation of bipotent signature with luminal progenitor status in FACS bulk data. A)** Boxplot of t-statistics of differential expression between the EPCAM+/CD49f+/ALDH+ cells (n=11) and all other luminal and putative luminal progenitors (n=27) from the FACS bulk data study of Shehata M et al, displaying these statistics for 17 genes upregulated in the bipotent single-cell cluster and which also map to the mammary stem-cell signature of Pece et al. **B)** As A), but now for all the 72-genes upregulated in the bipotent single-cell cluster.

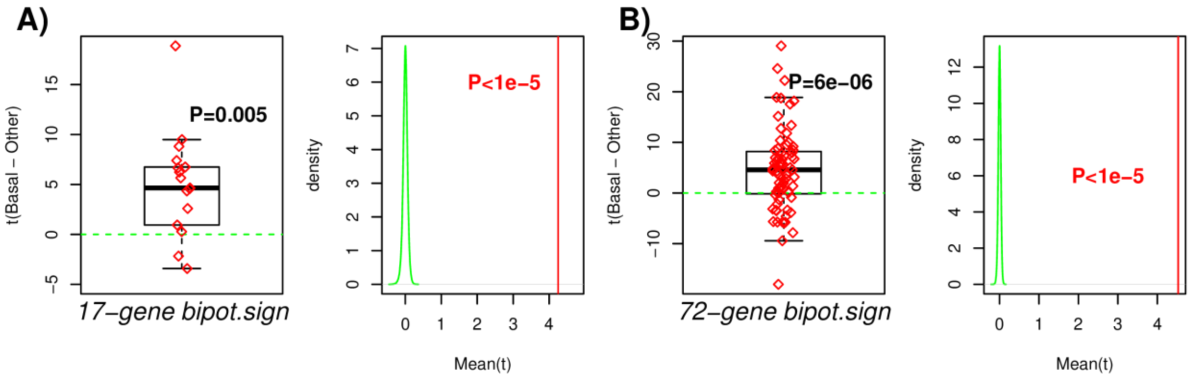


**fig.S12: Correlation of bipotent signature with basal breast cancer status in METABRIC. A)** Boxplot of t-statistics of differential expression between the basal breast cancers from METABRIC (n=329) and all other intrinsic subtypes (n=1645), displaying these statistics for the 17 genes upregulated in the bipotent single-cell cluster and which also map to the mammary stem-cell signature of Pece et al. **B)** As A), but now for all the 72-genes upregulated in the bipotent single-cell cluster.
